# Biofortification of green seaweed *Ulva* with Vitamin B12 using *Lacticaseibacillus rhamnosus* and *Lactococcus lactis*

**DOI:** 10.1101/2025.10.14.682258

**Authors:** S Sanjay, Meghanath Prabhu

## Abstract

One of the main causes of Vitamin B12 deficiency is insufficient dietary intake, especially among vegetarians and vegans. Seaweeds are emerging as sustainable biomass for producing value added products, including nutritionally enriched supplements. This study investigates the biofortification of green seaweed *Ulva* sp. with Vitamin B12 via co-fermentation using *Lacticaseibacillus rhamnosus* and *Lactococcus lactis*, aiming to develop a probiotic supplement. *Ulva* sp. was harvested from Goan shores, and the bacterial strains were obtained from cheese and cultivated for use in fermentation. The rationale behind this study was to integrate the de novo synthesis of Vitamin B12 with the conversion of Vitamin B2 into 5,6-dimethylbenzimidazole (DMB), leading to higher production of Vitamin B12, with the BluB enzyme serving as the key connecting factor. Vitamin B12 in fermented *Ulva* sp. was quantified using HPLC with a C18 column at 361 nm, with acetonitrile and water as the mobile phase. The highest concentration was observed after 72 hours of fermentation, reaching 19.6 μg/mL, approximately five times higher than the control. These findings highlight the potential of *Ulva* sp. as a substrate for Vitamin B12 production through bacterial fermentation and its promising application as a probiotic supplement.

## 1. Introduction

Vitamin B12 (cobalamin) is an essential water-soluble vitamin involved in various key physiological functions, including red blood cell formation, DNA synthesis and more [1]. It is found naturally in various animal-based foods, including meat, fish, eggs, and dairy-based products including curd (yogurt), cheese, etc., and is quite absent in plant-based food sources [2]. Vitamin deficiencies, including a lack of vitamin B12, have recently been reported to affect people of various ages including pregnant and lactating women worldwide. Strict vegetarians are even more at play on the line for the need for vitamin B12. Various forms of the intricate vitamin B12 exist, and each one has a unique chemical makeup and biological activity. The most prevalent kinds are methylcobalamin, cyanocobalamin, hydroxocobalamin, and adenosylcobalamin. These vitamin B12 forms each function with a specific purpose with specific importance in the human body. Approximately 62% of vegetarians experience vitamin B12 deficiencies, compared to 25 to 86%, 21 to 41%, and 11 to 90% of children, adolescents, and the elderly. Adults recommended daily intake (RDI) is 2.4 mcg of vitamin B12. Pregnant or nursing women may need higher doses of vitamin B12, with an RDI of 2.6–2.8 mcg daily. The RDIs for infants and young children range from 0.4 mcg to 1.8 mcg per day [3]. Vitamin B12 is also essential for maintaining optimal cognitive function and preventing cognitive decline. Vitamin B12 and folate work together to produce healthy red blood cells. Megaloblastic anemia, caused by vitamin B12 deficiency, causes enlarged red blood cells and results in fatigue and reduced oxygen transport [4]. The increasing demand for natural and safe food products has resulted in an increased interest in the use of functional foods that supply added health benefits. Vitamin B12 is produced only by certain bacteria and is not available from plant-based sources. Therefore, the need for sustainable and efficient methods to produce Vitamin B12 is of high importance [5]. Seaweeds are a diverse group of multicellular marine macroalgae found in various marine environments across the globe. They are classified into three briny groups based on their pigment composition - red, green, and brown algae [6]. Seaweed can grow in a broad range of environmental conditions, from shallow waters that are warm with excess nutrients to those that are cold with fewer nutrients.

Seaweeds have been used for centuries for their nutritional and curative benefits, such as boosting the immune system, lowering blood pressure, and promoting healthy skin.

Seaweeds are the substantive sources of food, medicine, and industrial products and are widely used in various industries, including biotechnology, agriculture, and cosmetics [7]. They are also rich in vitamins and minerals and are a promising source of bioactive substances with the potential to be leveraged in the creation of novel medications, cosmetics, and solid food additives [8]. Seaweeds are naturally known to contain small amounts of Vitamin B12 (54.5 to 58.6 micrograms per 100 g dry mass) and are a promising substrate for the production of Vitamin B12 by microbial fermentation. *Lactobacillus* and *Propionibacterium* species are lactic acid bacteria that are unremarkably used in the fermentation of dairy and other functional food products [9]. These bacteria can synthesize Vitamin B12 for their biochemical needs and are known to metabolize a range of substrates including those derived from seaweeds. Commercial uses of these bacteria include the production and selling of vitamin B12 supplements. A de novo pathway involving a series of enzymatic reactions using amino acids such as glutamate as the initial source is used by bacteria, including *Lactobacillus* sp., to synthesize vitamin B12. Conversely, *Propionibacterium* sp., synthesize vitamin B12 by converting vitamin B2 to DMB (5, 6-dimethyl benzimidazole), which is then converted to vitamin B12. Both of the aforementioned types of bacteria are used in the strategy for the enhanced synthesis of vitamin B12, with BluB enzyme (5, 6-dimethyl benzimidazole synthase) serving as a connecting link between both pathways. *Lactobacillus* sp. lacks the gene factor encoding the BluB enzyme, which is necessary to transition riboflavin (vitamin B2) to cyanocobalamin (vitamin B12), but *Propionibacterium* sp. type bacteria do [10]. In addition to the synthesis of vitamin B12, other biomolecules such as vitamin B2 are also produced. Therefore, in summation to the de novo pathway for synthesizing vitamin B12, bacteria with genes for the BluB enzyme also carry out the BluB enzyme’s transition of vitamin B2 to vitamin B12 [11]. Vitamin B12 synthesis is anticipated to occur as a result of this fortification. Studies on this topic have shown that the co-fermentation of the antecedently stated bacterial types can make more vitamin B12 than either type of fermentation alone [12].

Green seaweed, *Ulva* sp., was selected as a substrate because it is edible, promptly available, and has high carbohydrate and protein content. *Ulva* has been found to hold a high proportion of essential amino acids, including lysine, leucine, and valine. It also contains non-essential amino acids, such as aspartic acid, glutamic acid, glutamate, and proline.

The amino group acid content of *Ulva* may mostly depend on the season, location of the seaweed, and environmental conditions. Nevertheless, it is considered to be one of the most nutritionally beneficial seaweeds due to its high protein and amino acid content [13]. The overarching aim of this research study was to look into the co-fermentation of seaweed using *Lactobacillus* and *Propionibacterium* species for the enriched synthesis of Vitamin B12. For this purpose, samples of *Ulva* sp. were collected, identified and characterized, followed by fermentation of *Ulva* sp. for the production of Vitamin B12. The overall reaction pathway involved is shown in Fig.1.

**Fig. 1.**
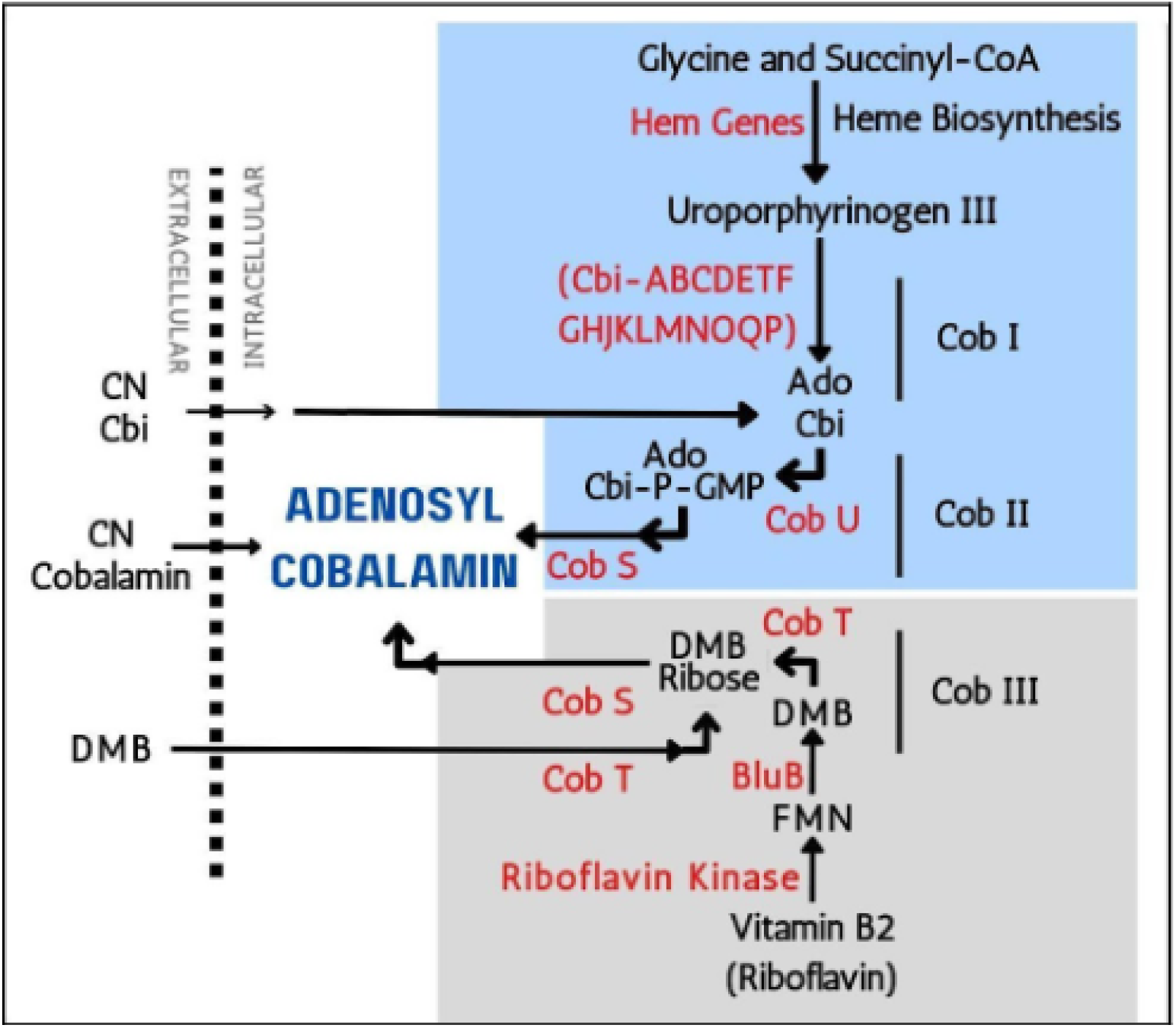
Schematic representation of all the reactions involved in the synthesis of Vitamin B12 via Co-fermentation of *Ulva* sp.

## 2. Materials and methods

### 2.1: Collection, identification and storage of green seaweed *Ulva* sp

*Ulva* sp. was collected from the shores of Vagator Beach in Goa. The green seaweed was manually harvested using the common hand harvesting technique by wading through shallow water and scooping it out of the rock surfaces to which it was attached. The seaweed was identified based on morphological characteristics. In order to get rid of any sand, dirt, or impurities, the collected seaweed was thoroughly cleaned and rinsed with filtered seawater. To avoid moisture loss and freezer burn, it was then drained and packaged in airtight plastic bags and stored in -20°C until further use [14].

### 2.2: Isolation and identification of *Lactobacillus* and *Propionibacterium* species

#### 2.2.1: Isolation of *Lactobacillus* species bacteria

Numerous dairy products, including yogurt, cheese, and kefir drinks, contain *Lactobacillus* species [15]. Cheddar cheese was thereby used in *Lactobacillus* isolation. The MRS agar is a specific media for *Lactobacillus* species. MRS media is an exclusive culture medium made to encourage *Lactobacilli’s* prolific development for laboratory research. For the purpose of this work, selective media, *Lactobacillus* MRS Agar (SRL cat. no. “79562”) was used. A sample of 1 gram of cheddar cheese was taken and placed in a sterile test tube with 9 ml sterile 0.85% saline. It was then homogenized to create a smooth paste. In order to promote individual colony growth on petri dish media, the homogenized cheese sample was streaked using quadrant streaking method. The plates were incubated for 48-72 hours at 37 °C and were then examined for the development of bacterial colonies. The bacterial colonies were picked out based on the morphology and further purification by streaking on MRS agar plates.

The *Propionibacterium* species were isolated using the same procedure as described above, with the exception of the culture medium used. The presence of *Propionibacterium freudenreichii*, a probiotic grade *Propionibacterium* species, was expected in milk-based products such as cheese. Therefore, the same cheddar cheese sample was used for the isolation. A selective medium that promotes growth of *Propionibacterium* species, while inhibiting that of other bacterial species, is yeast extract lactate agar with cadmium chloride. The preparation of this medium involved the incorporation of cadmium chloride into yeast extract lactate agar. After the growth was obtained, further evaluation of the pure isolates was carried out to determine the *Propionibacterium* species.

### 2.3: Identification of the isolated bacterial cultures

#### 2.3.1: Gram Staining

The morphology of the bacterial isolates was identified by Gram staining method. The bacterial smear after staining was observed using a light microscope and 100x objective lens with oil immersion [16].

#### 2.3.2: Molecular identification using 16s ribosomal RNA gene sequencing

Freshly streaked pure bacterial isolate was sequenced using the Hi-Gx360 Service of 16s ribosomal RNA sequencing. The sequencing was carried out at HiMedia Laboratories Pvt. Ltd. of Thane, Maharashtra, India. The sequence obtained was subjected to BLAST in the NCBI database for identification and phylogenetic tree based on the NJ (Neighbor-joining) algorithm [17].

### 2.4: Fermentation of *Ulva* sp. for the production of Vitamin B12

#### 2.4.1: Preparation of inoculum

A loopful of both the bacterial cultures were separately inoculated in yeast extract lactate broth and incubated at 37 °C. Assuming a linear relationship between OD 600 and bacterial cell density, an OD 600 value of 1 typically corresponds to a bacterial cell density in a bacterial broth culture of about 1x10^8^ cells/mL. In order for both of the chosen bacterial cultures, which were inoculated in broth, to reach the 1x10^8^ cell count, they were incubated for a period of time of approximately 16 to 20 hours. The incubation was ceased once the cultures’ respective OD at 600 nm values crossed the OD value of one [18].

#### 2.4.2: Seaweed Processing

The seaweed biomass (wet) and water were mixed in a 1:4 ratio and homogenized to obtain paste. Here, the homogenized mixture serves as the sample of the substrate, and 5 mL of the sample was transferred to four sterile 20 mL flasks. Given that vitamin B12 is photosensitive and thus susceptible to deterioration, the flasks were covered with aluminum foil.

#### 2.4.3: Fermentation of the seaweed

The “Test” flasks containing seaweed samples as substrate were inoculated with both of the isolated bacterial cultures from the broth. At the rate of 5%, 250 μL of each bacterial culture from the broth was transferred to the two ‘Test’ flasks for the co-fermentation process. Five hundred μL of sterile distilled water was added to the ‘Control’ flasks.

The flasks were placed in a shaker incubator set to 37 °C and 100 rpm. Under the aforementioned conditions, incubation was carried out continuously for 5 days.

### 2.5: Quantification of Vitamin B12 in the fermented seaweed

#### 2.5.1: Collection and Storage

Five hundred μL of the fermented seaweed sample from each flask was collected once every 24 hours for five days in 1 mL vials covered with aluminum foil, and it was then stored at -20°C until used for the extraction of vitamin B12.

#### 2.5.2: Extraction Vit B12 from the fermented seaweed sample

Hundred μL of fermented *Ulva* sp. samples from days 1, 3, and 5 of both the “Test” and “Control” flasks were pipetted out. The samples were then transferred to 2 mL centrifuge tubes (wrapped in aluminum foil) to which 100 μL of 0.05 mol/L acetate buffer (pH 4.8) was added. Next, 20 mg of potassium ferrocyanide was added, and the total volume was then brought to 1 ml by using distilled water. After that, the samples were kept in a boiling water bath for 30 minutes followed by centrifugation for 10 minutes at 10000 g, and the supernatant was collected for further Vitamin B12 analysis using HPLC [19].

#### 2.5.3: HPLC analysis for the quantification of total cyanocobalamin

The HPLC analysis of the extract was carried out using “Shimadzu LCMS-2020” with a 150 mm x 4.6 mm C18 column and its own software for data integration. A gradient of acetonitrile and water was used as a mobile phase solvent at a flow rate of 1 mL/min. A 20 μL extract sample was injected via injection port. The integrated data was then analyzed for the quantification of cyanocobalamin [20]. The peak areas of the respective were marked down and used for further analysis. The calibration curve was plotted using Microsoft Excel with the standard concentration vs peak area. And the equation of the straight was calculated to be Y = 1441.7X + 7607.8 with a Coefficient of determination, R^2^ = 0.971. The concentration of all the individual samples was calculated using this equation substituting ‘Y’ with the value of the sample’s peak area obtained. Calibration curves were constructed in Matplotlib using standards of known concentrations plotted against their respective peak areas. The analytical script, standard concentrations, and resulting calibration equations are detailed in the supplementary materials. Concentrations of all samples were subsequently determined based on the derived calibration equations and are reported in Table 1 and Table 2.

**Table 1.** (A) Peak area and the calculated concentration for Control. (B) The mean calculated concentration over 3 days.

| Control | Peak area | Concentration<br>mcg/mL |
| --- | --- | --- |
| C.1.0 | 7957 | 0.24 |
| C.1.3 | 17300 | 6.72 |
| C.1.5 | 17698 | 7.0 |
| C.2.0 | 11034 | 2.37 |
| C.2.3 | 9687 | 1.44 |
| C.2.5 | 9346 | 1.20 |

(B)
| Days | Concentration<br>mcg/mL |
| --- | --- |
| C.0 | 1.31 |
| C.3 | 4.08 |
| C.5 | 4.10 |

**Table 2.** (A) Peak area and the calculated concentration for the Test. (B) The mean calculated concentration over 3 days.

| Test | Peak Area | Concentration<br>mcg/mL |
| --- | --- | --- |
| T.1.0 | 11133 | 2.44 |
| T.1.3 | 31091 | 16.28 |
| T.1.5 | 27481 | 13.78 |
| T.2.0 | 17557 | 6.90 |
| T.2.3 | 40843 | 23.05 |
| T.2.5 | 25267 | 12.24 |

(B)
| Days | Concentration<br>mcg/mL |
| --- | --- |
| T.0 | 4.67 |
| T.3 | 19.67 |
| T.5 | 13.02 |

## 3. Results and discussion

### 3.1 Identification of collected seaweed

*Ulva* species can be identified at the macroscopic level by their distinctive thin, flat, bright green thallus, which often appears translucent when submerged in water. These algae typically exhibit a broad, blade-like morphology with irregular lobes and undulations along the margins, giving them a visually unique appearance [21]. Based on these morphological criteria the collected seaweed was identified to be of *Ulva* sp.

### 3.2: Identification of the isolated Bacteria

#### 3.2.1: Isolate 1

The isolate obtained on the *Lactobacillus* MRS agar was expected to be of any *Lactobacillus* species. The isolate was found to be Gram Positive bacteria (supplementary **Fig. 1**). Therefore, it was selected for further procedures.

#### 3.2.2: 16s rRNA Sequencing

A reliable method for identifying and classifying bacteria according to their genetic makeup is 16s ribosomal RNA gene sequencing. The technique is based on the sequencing of a conserved region of the 16s ribosomal RNA gene of bacteria, which experiences minute changes over time and enables the determination of taxonomic relationships among various bacterial strains. In order to produce accurate data, the protocol calls for DNA extraction, PCR amplification of the 16s ribosomal RNA gene, and DNA sequencing of the amplicons. More accurate bacterial identification can be achieved by comparing the generated sequence with reference databases and using bioinformatics tools to find the closest-matching bacterial sequence. The freshly streaked isolates were sent for 16s rRNA sequencing as mentioned in the section “2.2.1”. The sequence was obtained for the sequencing process involved with primer “27R” and the BLAST results showed 95.39% similarity with ‘*Lacticaseibacillus rhamnosus*’. And the culture was identified as similar to that of the bacterial species *Lactobacillus rhamnosus*.

The Neighbor-Joining based phylogenetic tree (**Fig. 2**) was constructed for result interpretation. The phylogenetic tree constructed depicted the relatedness of the isolate with that of others and based on the results of the phylogenetic tree analysis for 16S rRNA gene sequence that has 95.3% similarity to the bacterial species *Lacticaseibacillus rhamnosus*, the sequence is likely to belong to the same genus as *L. rhamnosus*, which was previously classified as *Lactobacillus*.

**Fig. 2.**
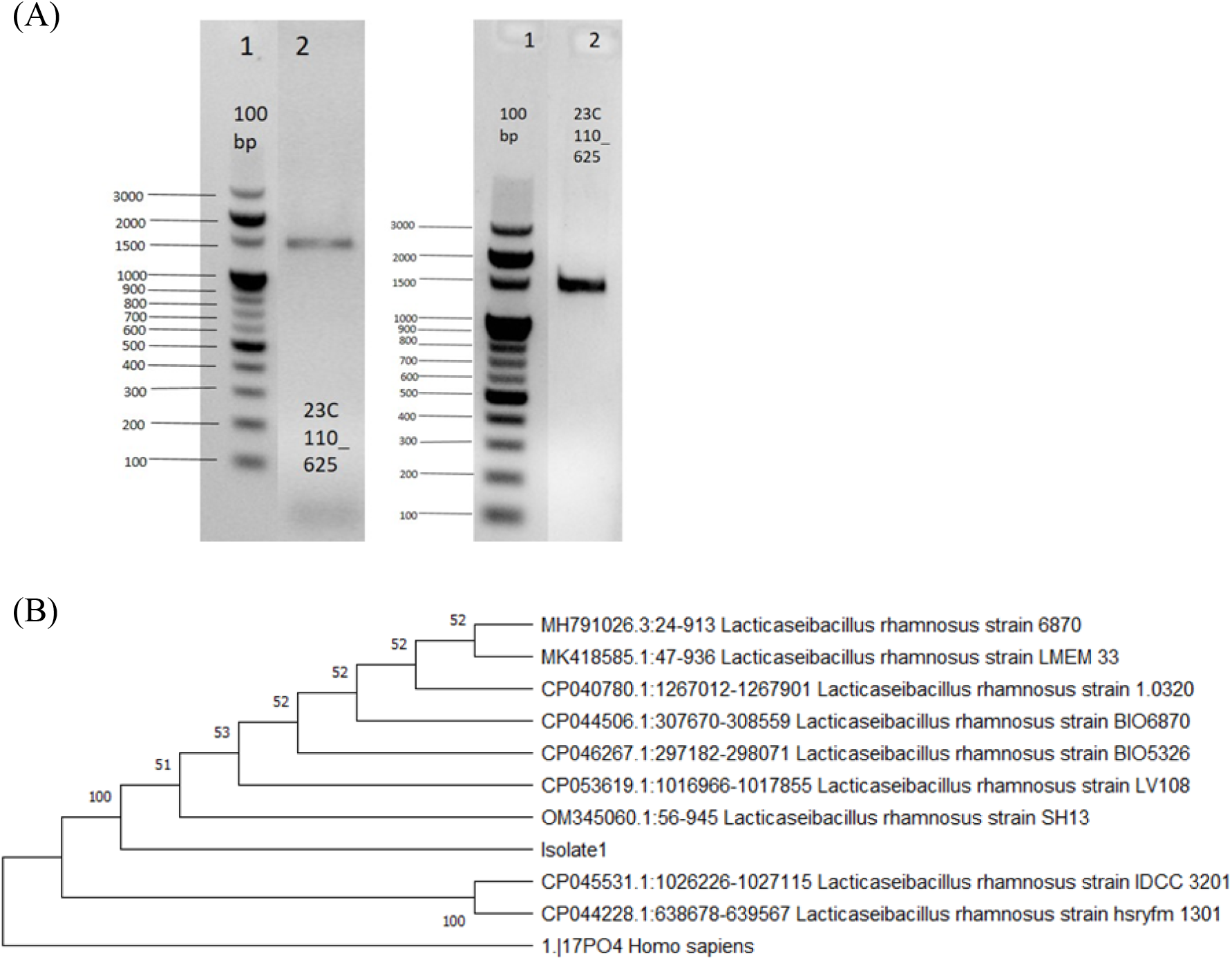
(A) Resulted PCR Gel-Image of the Isolate 1 confirming the isolation of 16S rRNA gene of 1550 base pairs with that of reference [22]. (B) The constructed phylogenetic tree using the Neighbour Joining method. With the incidence of 95.3% similarity, it can be grouped under the species of *Lacticaseibacillus rhamnosus*.

The 16s rRNA sequence of the isolated ‘*Lacticaseibacillus rhamnosus*’ is available on the GenBank database with Id - OR294944.

#### 3.2.3: Isolate 2

The first isolate which was cultivated on the Yeast Extract Lactate agar and expected to be of any *Propionibacterium* species underwent a preliminary analysis involving gram staining followed by 16S rRNA sequencing. The results were interpreted.

#### 3.2.4: Gram Staining

The gram staining technique was carried out and the result was found to be Gram Positive Bacteria (supplementary **Fig**. 2). Therefore, the isolate was selected for further procedures.

#### 3.1.5: 16s rRNA Sequencing

The freshly streaked isolates were sent for 16s rRNA sequencing as mentioned in the section “3.2.3.2”. The sequence was obtained for the sequencing process involved with primer “907R” and the BLAST results showed 100% similarity with the species ‘*Lactococcus lactis*’. And therefore, probably predicted in high confidence to be the bacterial species *Lactococcus lactis*.

The Neighbor-Joining based phylogenetic tree was constructed for result interpretation. The result of phylogenetic tree analysis for a 16S rRNA gene sequence having 100% similarity with the bacterial species *Lactococcus lactis* indicates that the sequence belongs to the same species as *Lactococcus lactis* (Fig. 3). The 16s rRNA sequence of the isolated ‘*Lactococcus lactis*’ is available on the GenBank database with Id - OR294964.

**Fig. 3.**
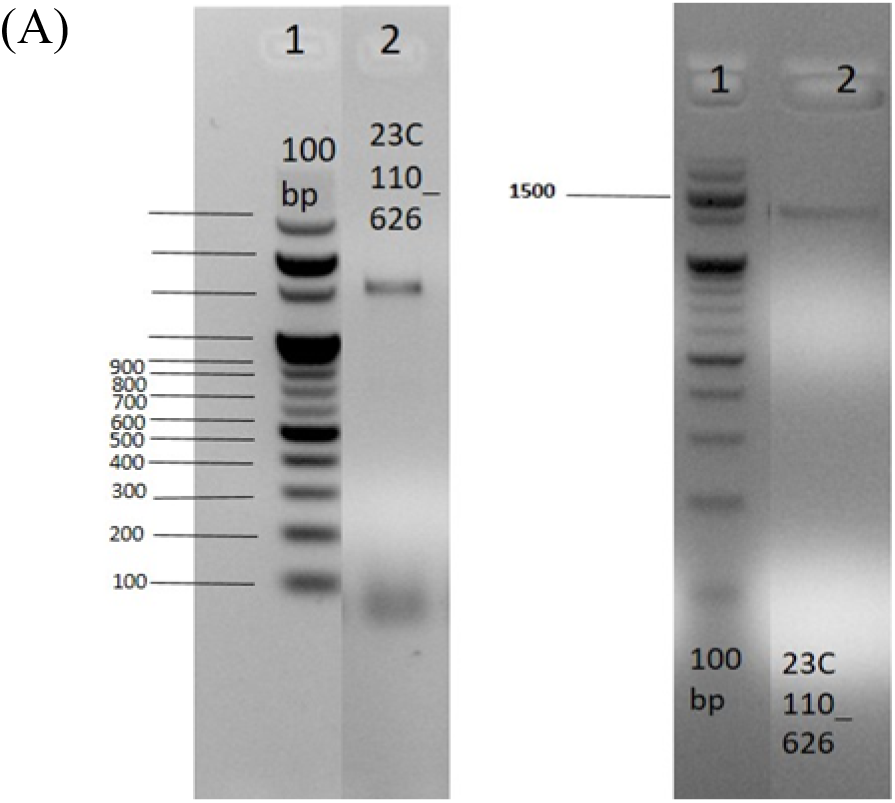
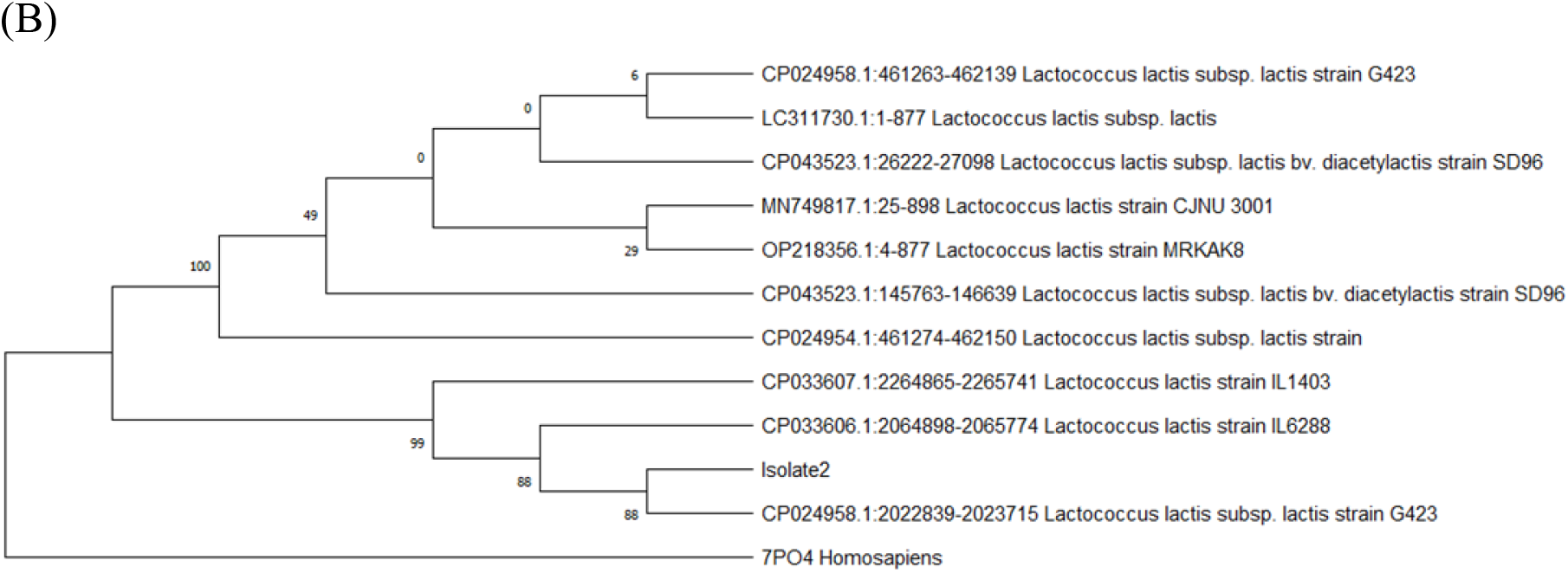
(A) Resulted PCR Gel-Image of the Isolate 1 confirming the isolation of 16S rRNA gene of 1550 base pairs with that of reference, (B) The constructed phylogenetic tree using the Neighbour Joining method. With the incidence of 100% similarity, it can be grouped under the species of *Lactococcus lactis*.

Despite the fact that we isolated the *Propionibacterium* species using specialized conditions with high metal concentrations, the isolate has been identified as *Lactococcus lactis*. As a result, research demonstrates that *Lactococcus lactis* may thrive in heavy metal media, however many other bacteria cannot tolerate and grow. The reason for choosing *Propionibacterium* species was the ability to possess the ‘BluB’ gene, which is essential for the synthesis of Vitamin B12 by converting FMN to DMB. Therefore, a NCBI database search was conducted to see if the ‘BluB’ gene has been reported in *Lactococcus lactis*, and the findings were positive. Multiple entries in UniProt for the presence of the ‘BluB’ gene in ‘*Lactococcus lactis*’ are reported and one such entry for confirmation is with an ID - A0A0H1RRD1.

### 3.3: HPLC Analysis for the Quantification of Cyanocobalamin

The other forms of cobalamin, except cyanocobalamin, eventually degrade because they are unstable during extraction and analysis. Therefore, the other forms of cobalamin are converted into the stable cyanocobalamin form, which makes it simple to measure total cobalamin, with the addition of a cyanide compound, potassium ferrocyanide. HPLC analysis using the C18 column proved to be a reliable and accurate method for the analysis of cyanocobalamin in vitamin B12 samples.

The combination of the C18 column and gradient system produced the high-resolution separation of the analytes with minimal interference from other compounds. The method’s excellent detection and quantification limits allowed for an accurate and precise analysis of cyanocobalamin in the sample. The method of preparation was carried out using one of the standards i.e., 1 mcg/mL standard and the cyanocobalamin’s peak was seen and noted around the retention time of 6.6 mins. For all the standards and samples the peaks were observed in the range of 6.4 to 6.7 mins. The mean was taken for the calculated concentration of each ‘Control’ and ‘Test’ respectively at Day 0, 3, and 5. The ‘C.1 and C.2’ represent the Control in duplicates. In the same way the ‘T.1 and T.2’ denote the test in duplicates.

The amount of cyanocobalamin in the ‘Control’ sample on day 0, day 3 and, and day 5 day was determined to be 1.3, 4.08, and 4.1 mcg/mL respectively. The amount of cyanocobalamin in the ‘Control’ did not have much deviations in the value over a period of time. This confirms the fact that seaweed does not have the capability to produce Vitamin B12 on its own. The amount of cyanocobalamin in the ‘Test’ sample (fermented seaweed) on day 0 day 3 and day 5 was determined to be 4.6, 19.6, and 13.01 mcg/mL (**Fig. 4**).

**Fig. 4.**
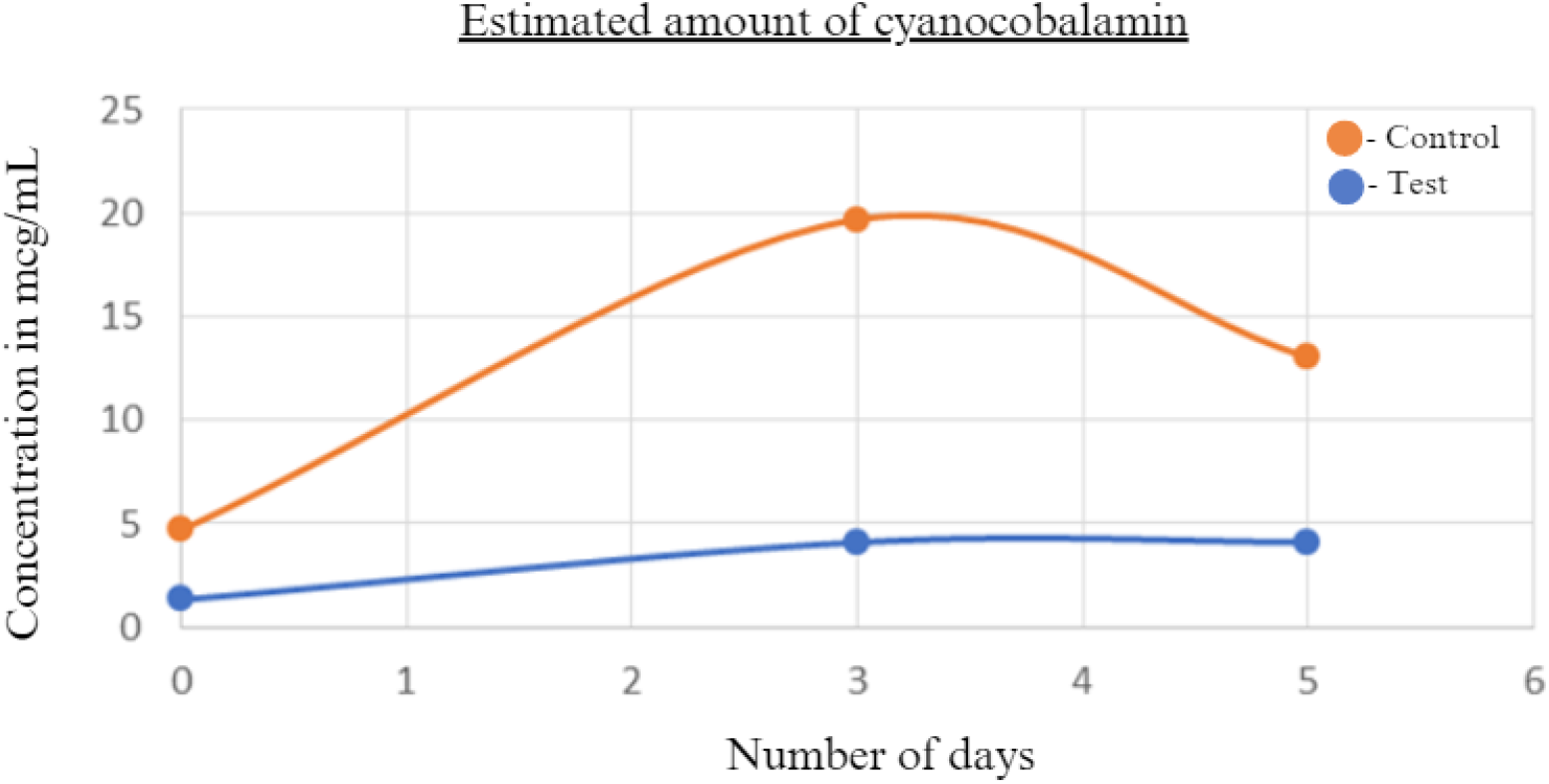
Graphical representation depicting the comparative values of the amount of Vitamin B12 produced on 0^th^, 3^rd^ and 5^th^ day.

This depicts the fortified production of Vitamin B12 by the probiotic bacteria used on the seaweed as substrate. The concentration of cyanocobalamin increases from the 0th day to the 3rd day and gradually decreases on the 5th day may be due to the fact that the bacteria utilize the produced Vitamin B12 for its proliferation. Therefore, it can be concluded that the optimum time for the maximum concentration of Vitamin B12 (19.6 mcg/mL), is around 72 hours which is during the 3rd day. A similar experiment has been carried out using coconut milk as a substrate, fermented by *Lactobacillus casei* L4, and the highest concentration of Vitamin B12 produced was 1.47 mcg/mL [23].

## 4. Conclusion

This research was intended to make a commercial product as a fusion of probiotics and products of nutrient supplements into one. The co-fermentation of seaweed as a substrate for the production of Vitamin B12 using *Lacticaseibacillus rhamnosus* and *Lactococcus lactis* proved to be successful. The concentration of Vitamin B12 was 19.6 μg/ml in the fermented seaweed (380.4 %higher compared to control) at the optimum period i.e., 72 hours.

This indicates that this method can be employed as a feasible and cost-effective way for the production of Vitamin B12 as a probiotic supplement. The use of seaweed as a substrate for fermentation presents a promising alternative to conventional methods that commonly use animal-derived products. The results of this study are particularly significant in light of current concerns about the availability and sustainability of Vitamin B12 sources. As this vitamin is essential for human health, particularly for those following a vegan or vegetarian diet, developing new and sustainable methods for its production is crucial.

This fermentation product can be used as a probiotic supplement with a fortified amount of Vitamin B12 which can meet the quantity of daily requirement. Overall, this research provides important insights into the potential use of co-fermentation with seaweed as a means of producing Vitamin B12. Further studies are required that can help to optimize the production process, make the product better for consumable grade with added flavors, and determine its commercial feasibility, paving the way for a more sustainable and accessible source of this crucial vitamin.

## Acknowledgements

This project was supported by funds provided by the Department of Biotechnology (DBT), India to Discipline of Biotechnology, School of Biological Sciences and Biotechnology, Goa University, as a part of the post-graduation course in Marine Biotechnology.

## Notes

### Competing Interest Statement

The authors have declared no competing interest.

