## Supplemental Files for "Biofortification of green seaweed *Ulva* with Vitamin B12 using *Lacticaseibacillus rhamnosus* and *Lactococcus lactis*"

Supplementary **Fig. 1.** The bacteria retain crystal violet and therefore appear purple, confirming that it is Gram-Positive. And the rod shape of the bacteria supports it possibly could be of *Lactobacillus* species.

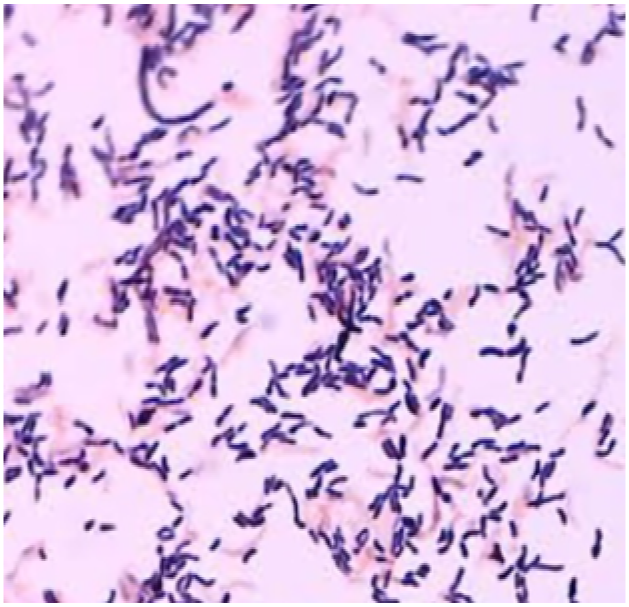

Supplementary **Fig. 2.** The bacteria retain crystal violet and therefore appear purple, confirming that it is Gram-Positive with short- rod morphology.

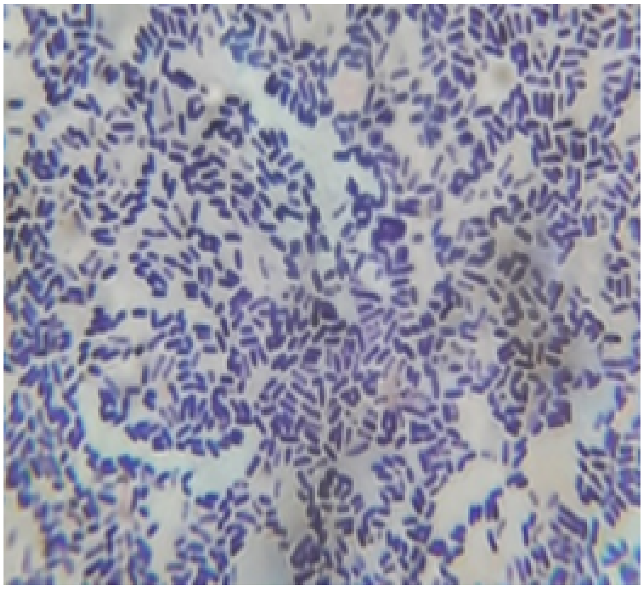

**Table. 1.** Standard concentration and its respective peak area in HPLC analysis. S1 to S6 represent the standards starting from 0.0001 mcg to 10 mcg.

| Standard Number | Concentration (mcg/mL) | Peak Area (a.u.) |
|---|---|---|
| S1 | 0.0001 | 7402 |
| S2 | 0.001 | 7115 |
| S3 | 0.01 | 8056 |
| S4 | 0.1 | 6479 |
| S5 | 1 | 10746 |
| S6 | 10 | 21867 |

**Fig. 3.** Standard Calibration Curve.

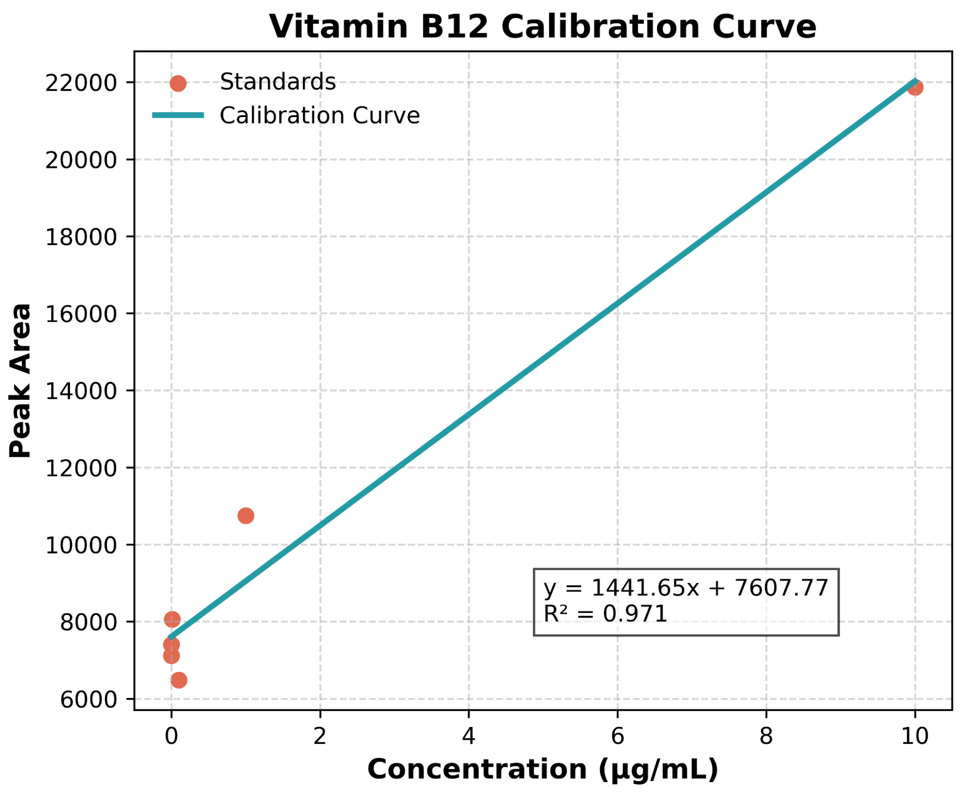

Analytical Script using Matplotlib:

```
import numpy as np
import matplotlib.pyplot as plt
from sklearn.linear_model import LinearRegression
# Data
concentration = np.array([0.0001, 0.001, 0.01, 0.1, 1.0, 10.0]).reshape(-1, 1)
peak_area = np.array([7402, 7115, 8056, 6479, 10746, 21867])
# Fit regression
model = LinearRegression()
model.fit(concentration, peak_area)
slope = model.coef_[0]
intercept = model.intercept_
r2 = model.score(concentration, peak_area)
# Line for plotting
x_line = np.linspace(0, 10, 200).reshape(-1, 1)
y_line = model.predict(x_line)
# Plot
plt.figure(figsize=(6,5))
plt.scatter(concentration, peak_area, color="#E16A54", s=40, label="Standards")  # smaller points
plt.plot(x_line, y_line, color="#239BA7", linewidth=2.5, label="Calibration Curve")
# Labels & style
plt.xlabel("Concentration (µg/mL)", fontsize=12, weight="bold")
plt.ylabel("Peak Area", fontsize=12, weight="bold")
plt.title("Vitamin B12 Calibration Curve", fontsize=14, weight="bold")
plt.text(5, 8000, f"y = {slope:.2f}x + {intercept:.2f}\nR² = {r2:.3f}",
         fontsize=10, bbox=dict(facecolor='white', alpha=0.7))
plt.legend(frameon=False)
plt.grid(True, linestyle="--", alpha=0.5)
plt.tight_layout()

# Save figure
plt.savefig("calibration_curve.png", dpi=600)
plt.show()
```
